# Chikungunya virus ECSA lineage reintroduction in the northeasternmost region of Brazil

**DOI:** 10.1101/2021.01.09.426024

**Authors:** Joilson Xavier, Vagner Fonseca, Joao Felipe Bezerra, Manoella do Monte Alves, Maria Angélica Mares-Guia, Ingra Morales Claro, Ronaldo de Jesus, Talita Adelino, Emerson Araújo, Karina Ribeiro Leite Jardim Cavalcante, Stephane Tosta, Themis Rocha de Souza, Flavia Emanuelle Moreira da Cruz, Allison de Araújo Fabri, Elaine Cristina de Oliveira, Noely Fabiana Oliveira de Moura, Rodrigo Fabiano do Carmo Said, Carlos Frederico Campelo de Albuquerque, Vasco Azevedo, Tulio de Oliveira, Ana Maria Bispo de Filippis, Rivaldo Venâncio da Cunha, Kleber Giovanni Luz, Marta Giovanetti, Luiz Carlos Junior Alcantara

## Abstract

The Northeast region of Brazil registered the second highest incidence proportion of chikungunya fever in 2019. In that year an outbreak consisting of patients presented with febrile disease associated with joint pain were reported by the public primary health care service in the city of Natal, Rio Grande do Norte state, in March 2019. At first, the aetiological agent of the disease was undetermined. Since much is still unknown about chikungunya virus (CHIKV) genomic diversity and evolutionary history in this northeasternmost state, we used a combination of portable whole genome sequencing, molecular clock, and epidemiological analyses that revealed the re-introduction of the CHIKV East-Central-South-African (ECSA) lineage into Rio Grande do Norte. We estimated CHIKV ECSA lineage was first introduced into Rio Grande do Norte in early June 2014, while the 2019 outbreak clade diverged around April 2018 during a period of increased chikungunya incidence in the Southeast region, which might have acted as a source of virus dispersion towards the Northeast region. Together, these results confirm the ECSA lineage continues to spread across the country through interregional importation events likely mediated by human mobility.

**HIGHLIGHTS:** CHIKV ECSA lineage introduction into Rio Grande do Norte state, Northeast Brazil, was estimated to early June 2014

At least two CHIKV importation events occurred in Rio Grande do Norte state, Brazil

The 2019 chikungunya outbreak in Rio Grande do Norte was likely caused by a second event of CHIKV introduction imported from Rio de Janeiro state.

## INTRODUCTION

Since its emergence in East Africa, chikungunya virus (CHIKV), an alphavirus belonging to the family *Togaviridae*, has caused more than 70 epidemics around the world mainly in Southeast Asian and Latin American countries (Mascarenhas et al., 2018).

Northeast Brazil presented with the second-highest incidence (59.4 cases per 100,000 population) of chikungunya notified in 2019 in the country (Ministério da Saúde, 2020b). In that year, an outbreak of 13,713 chikungunya cases was notified in the state of Rio Grande do Norte, in the far northeastern tip of Brazil, where a previous outbreak of almost 25,000 probable cases were reported in 2016 (Ministério da Saúde, 2018). Despite the high number of cases, much remains unknown about CHIKV genomic diversity and the evolutionary history in this northeastern state. In order to get more insight regarding CHIKV regional dispersion dynamic, we generated eight CHIKV genomes from the 2019 outbreak registered in Natal, the capital city of Rio Grande do Norte, and provided a genomic epidemiological report of the virus circulating in that state.

## METHODS

An outbreak consisting of patients presented with febrile disease associated with joint pain were reported by the public primary health care service in the city of Natal, Rio Grande do Norte state, in March 2019. Since the disease aetiological agent was undetermined at first, serum samples were provided to the State Central Laboratory of Public Health Dr Almino Fernandes and then sent to Laboratory of Flavivirus at the Oswaldo Cruz Institute (IOC) from the Oswaldo Cruz Foundation (Fiocruz) to confirm laboratory diagnosis by RT-qPCR (Lanciotti et al., 2007). CHIKV-RNA positive samples were submitted to a sequencing protocol based on multiplex PCR tiling amplicon approach and MinION nanopore sequencing (Quick et al., 2017). Final consensus sequences were *de novo* assembled by Genome Detective (Vilsker et al., 2019). Our new sequences were aligned to other publicly available sequences retrieved from NCBI. We performed Maximum likelihood (ML) and Bayesian time-scaled phylogenetic analysis using a molecular clock approach (more details in Appendix). We used chikungunya notified cases data from the Brazil Ministry of Health National Reporting System (SINAN-Net) (Ministério da Saúde, 2020a) to calculate incidence and to plot time series charts.

## RESULTS

Laboratory diagnostics confirmed 13 samples were positive for CHIKV RNA with a mean Ct value of 28.15. We generated eight CHIKV genomes from these 13 samples collected on 23/March – 01/April in 2019 from patients with an average 42 years of age. We found five (38%) patients presented with fever associated with myalgia, arthralgia, and rash (see Table 1 and Fig 1A). The eight CHIKV genomes generated here have an average genome coverage of 93% (ranging from 92% to 94%) and they belonged to the East-Central-South-African (ECSA) lineage as indicated by our ML phylogeny (see Fig. S1 and Table S1).

**Fig. 1.**
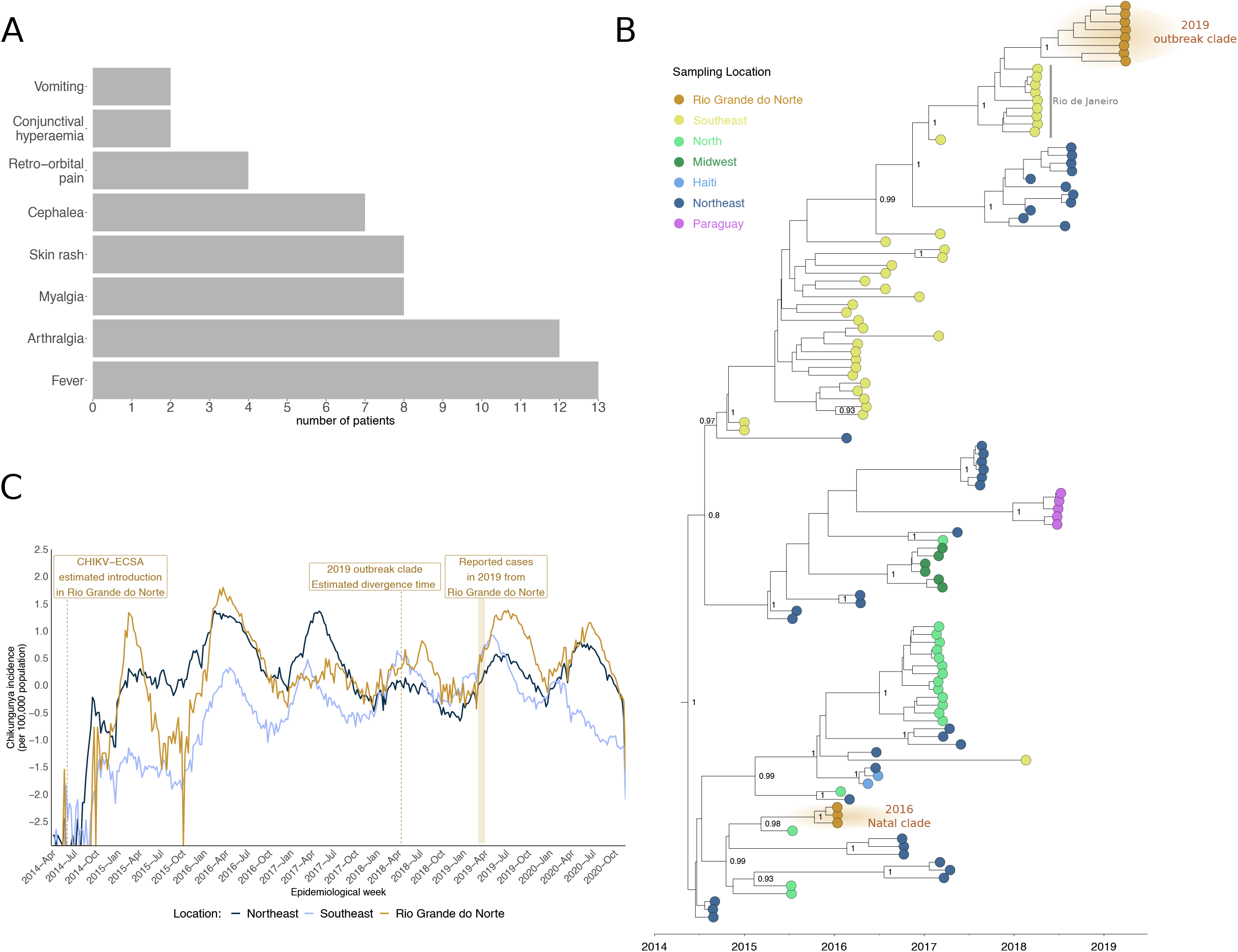
Symptoms frequency, epidemic curves and phylogenetic analysis from chikungunya outbreaks reported in Rio Grande do Norte state, northeast Brazil. (A) symptoms frequency among 13 patients with confirmed laboratory diagnosis of chikungunya infection. (B) Time-scaled maximum clade credibility tree of chikungunya virus East-Central-South African lineage in Brazil, including the 8 new genomes generated in this study. Tips are coloured according to sample source location. Values around nodes represent posterior probability support of the tree nodes inferred under Bayesian Evolutionary Analysis using a molecular clock approach. (C) Time-series plot of chikungunya incidence per 100,000 population calculated from notified cases from Rio Grande do Norte state, Northeast, and Southeast regions of Brazil. Y-axis values were log-transformed.

We inferred a time-measured phylogeny using a dataset comprising 110 Brazilian sequences plus two isolates from Haiti and five from Paraguay. Both our ML and Bayesian phylogenies showed the 2019 CHIKV isolates from Rio Grande do Norte formed a single well supported clade (hereinafter 2019 outbreak clade; posterior probability = 1.00), which was closely related to isolates from Rio de Janeiro (posterior probability = 1.00) (Fig. 1B). Interestingly, this 2019 outbreak clade did not cluster to the other three previously published isolates also from Natal, Rio Grande do Norte, sampled in January 2016 (hereinafter 2016 Natal clade; posterior probability = 1.00). We estimated the time of the most recent common ancestor (TMRCA) of the 2019 outbreak clade to be around mid-April 2018 (95% BCI: early December 2017 to late August 2018) (Fig. 1B and 1C). Moreover, the TMRCA of the CHIKV-ECSA lineage in Rio Grande do Norte state was estimated to be early June 2014 (95% BCI: late December 2013 to early October 2014).

Weekly reported incidences revealed three major outbreaks in Rio Grande do Norte state during early 2015, early 2016, and mid-2019. Smaller waves were observed throughout the years from 2017 to 2020, indicating persistence of the virus in the state through year-round transmission cycles (see Fig. 1C), which were also observed in the other regions of Brazil (Fig. S2).

## DISCUSSION

In this study we generated eight CHIKV genomes from symptomatic cases of chikungunya fever reported during an outbreak in March 2019 in Natal, the capital city of Rio Grande do Norte state, where 15,467 cases were notified in that same year. We found fever, arthralgia, myalgia and rash were the most common presenting symptoms, which are similar findings to those described in other studies on suspected cases during the 2016 chikungunya outbreak in Natal (Monteiro et al., 2020) and on chikungunya-associated neurological disease cases reported from 2014 to 2016 in another Brazilian northeastern state (Brito Ferreira et al., 2020).

Our phylogenetic analysis revealed the 2019 outbreak clade is distant related to other isolates from the previous 2016 outbreak in Rio Grande do Norte. This suggests at least two importation events of CHIKV-ECSA lineage occurred in Rio Grande do Norte, being the time of the first virus importation event estimated to be early June 2014. This estimate indicates the virus was circulating for several months before the first chikungunya cases were confirmed in 2015 in Rio Grande do Norte, including one first chikungunya-associated death, as indicated by available epidemiological surveillance data from the State Health Department (Secretaria da Saúde Pública, 2017).

From combined analysis of regional epidemic curves and time-scaled phylogeny, we hypothesize that the 2019 outbreak in Rio Grande do Norte was caused by a second event of CHIKV introduction imported from Rio de Janeiro. We estimate the divergence time of the 2019 outbreak clade to be mid-April 2018, which corresponds to a period of increased chikungunya incidence in the southeast region, where Rio de Janeiro state is located. This might be explained by virus interregional spread through people movement, as discussed elsewhere (Candido et al., 2020; Churakov et al., 2019).

Our study increases the number of CHIKV genomic sequences and consequently provides more information on evolution and genomic diversity of the ECSA lineage currently circulating in the Northeast region of Brazil, where this lineage was first introduced. Hence, continuing arbovirus genomic surveillance can contribute to a comprehensive understanding of CHIKV evolution and spread in the country, thus assisting epidemic control and prevention in at-risk populations.

## Ethics statement

This project was reviewed and approved by the Comissão Nacional de Ética em Pesquisa (CONEP) [National Research Ethics Committee], as part of the arboviral genomic surveillance efforts within the terms of Resolution 510/2016 of CONEP, by the Pan American Health Organization Ethics Review Committee (PAHOERC) (Ref. No. PAHO-2016-08-0029), and by the Oswaldo Cruz Foundation Ethics Committee (CAAE: 90249218.6.1001.5248).

## Data availability

Newly generated CHIKV sequences have been deposited in GenBank under accession numbers <u>**MW260512-MW260519**</u>.

---

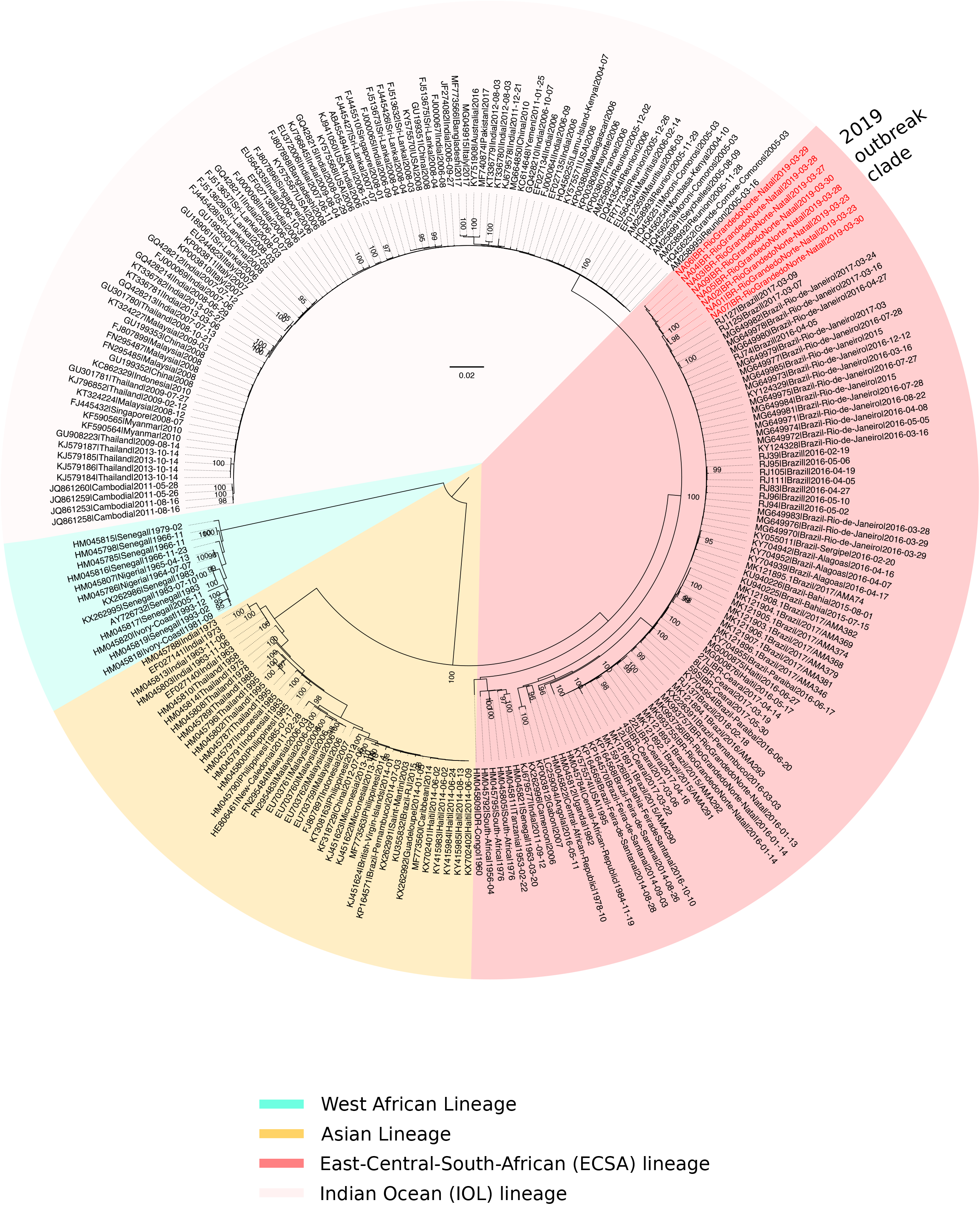

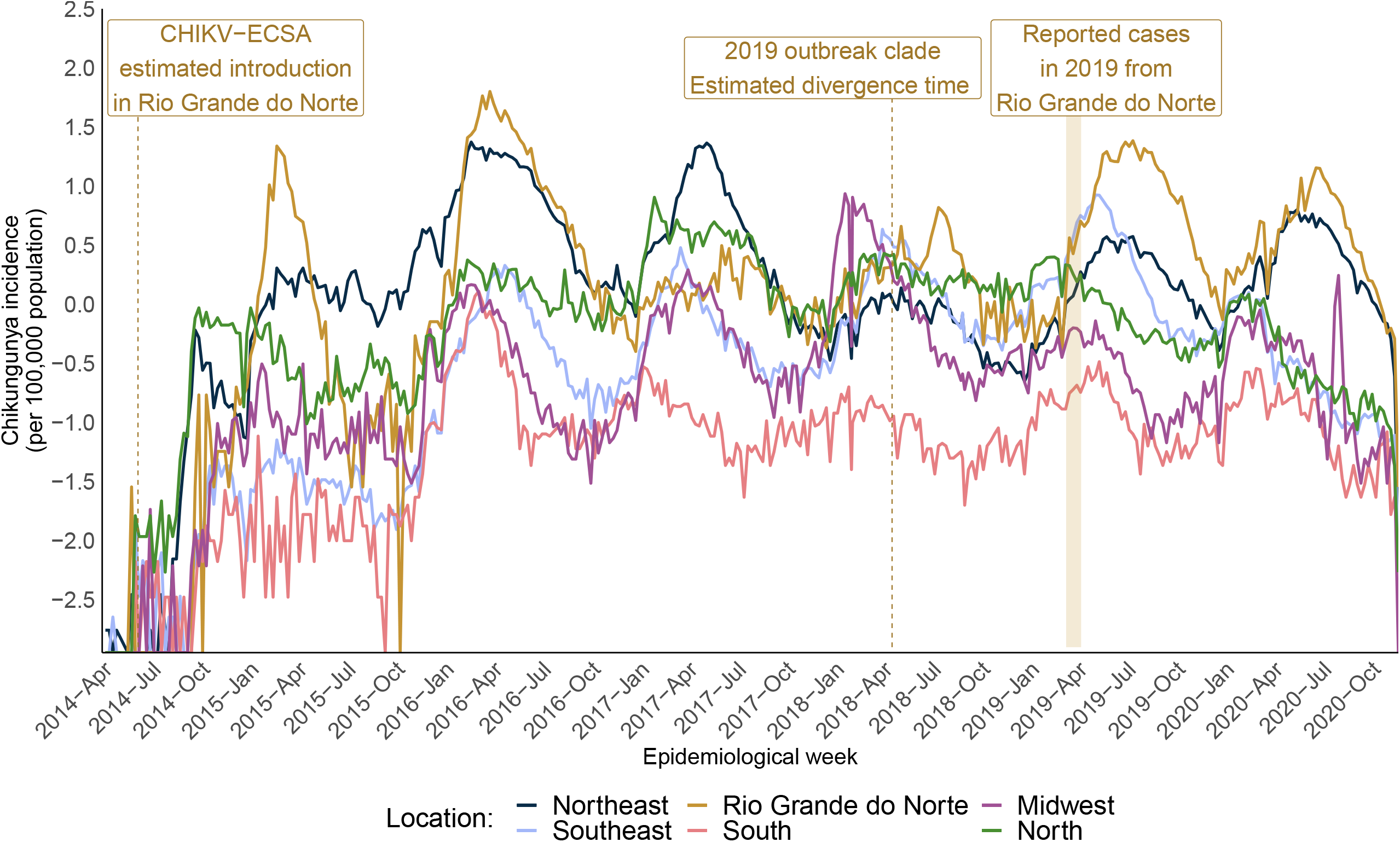

