## Supplementary files for "Chikungunya virus ECSA lineage reintroduction in the northeasternmost region of Brazil"

**METHODS**

**Sample collection and laboratory diagnosis**

Serum samples from individuals presented with febrile disease associated with joint pain were collected for molecular diagnostics performed at the State Central Laboratory of Public Health Dr Almino Fernandes, Rio Grande do Norte state, northeast Brazil. Then, to confirm chikungunya infection, those samples were sent to the Brazilian national reference laboratory for arbovirus diagnosis at the Laboratory of Flavivirus in the Oswaldo Cruz Institute (IOC) from the Oswaldo Cruz Foundation (Fiocruz). Serum samples were submitted to nucleic acid purification using the MagMax Viral RNA Mini kit (Thermo Fischer Scientific), following manufacturer’s recommendations. RT-qPCR tests were performed on 13 samples sent to Fiocruz, where chikungunya infection was confirmed (Lanciotti et al., 2007).

**Sequencing experiments**

Samples were submitted to cDNA synthesis and PCR, using a sequencing protocol based on multiplex PCR tiling amplicon approach design for MinION nanopore sequencing (Quick et al. 2017). DNA library preparation was performed on 8 selected samples (selection based on DNA concentration after clean-up being > 4ng/μL) using the Ligation Sequencing Kit (Oxford Nanopore Technologies). Individual samples were barcoded using the Native Barcoding Kit (NBD104, Oxford Nanopore Technologies, Oxford, UK). Sequencing library was loaded onto a R9.4 flow cell and data was collected for up to 48 sequencing hours.

**Generation of consensus sequences**

We used Guppy (https://github.com/nanoporetech) for basecalling of raw FAST5 files, while qcat (<https://github.com/nanoporetech/qcat>) was used for barcode demultiplexing. Consensus sequences were generated by *de novo* assembling using Genome Detective (<https://www.genomedetective.com/>) (Vilsker et al. 2019).

**Phylogenetic and Bayesian analysis**

The 8 new sequences reported in this study were initially submitted to a genotyping analysis using the phylogenetic arbovirus subtyping tool from Genome Detective (<http://genomedetective.com/app/typingtool/chikungunya>) (Fonseca et al. 2019). We used MAFFT (Katoh et al. 2005) to align these 8 new sequences to other 224 closely related CHIKV sequences publicly available in NCBI. We assessed phylogenetic signal on this dataset (n=232) using the likelihood mapping analysis implemented in IQ-TREE 2.1.1 software (Nguyen et al. 2015). We used IQ-TREE software to infer a Maximum Likelihood (ML) phylogeny under the GTR+F+I+G4 best fit substitution model identified by ModelFinder (Kalyaanamoorthy et al. 2017).

From the ML tree inferred from the dataset we constructed a small sub-dataset (n= 117) containing mainly CHIKV ECSA Brazilian sequences according to the Brazilian clade observed in the ML tree. We used TempEst (Rambaut et al. 2016) to assess the presence of temporal signal. To infer a time-scaled phylogeny from the sub-dataset, we used BEAST v1.10.4 (Suchard et al. 2018) to perform Bayesian phylogenetic analysis under GTR+G4 codon partition (CP)1+2,3 model. We combined sextuplicate independent runs of 50 million states each. For these runs, we used the uncorrelated relaxed molecular clock and the Bayesian skygrid population model (Gill et al. 2013). For comparison, we also used the strict molecular clock with the Bayesian skygrid population model. Maximum clade trees were summarized from the MCMC samples using TreeAnnotator after discarding 10% as burn-in.

**Epidemic curves from Chikungunya cases reported in Brazil**

We used chikungunya notified cases data from the Brazil Ministry of Health National Reporting System (SINAN-Net) (Ministério da Saúde, 2020) to calculate incidence and to plot time series charts.

**SUPPLEMENTARY FIGURES AND TABLES**

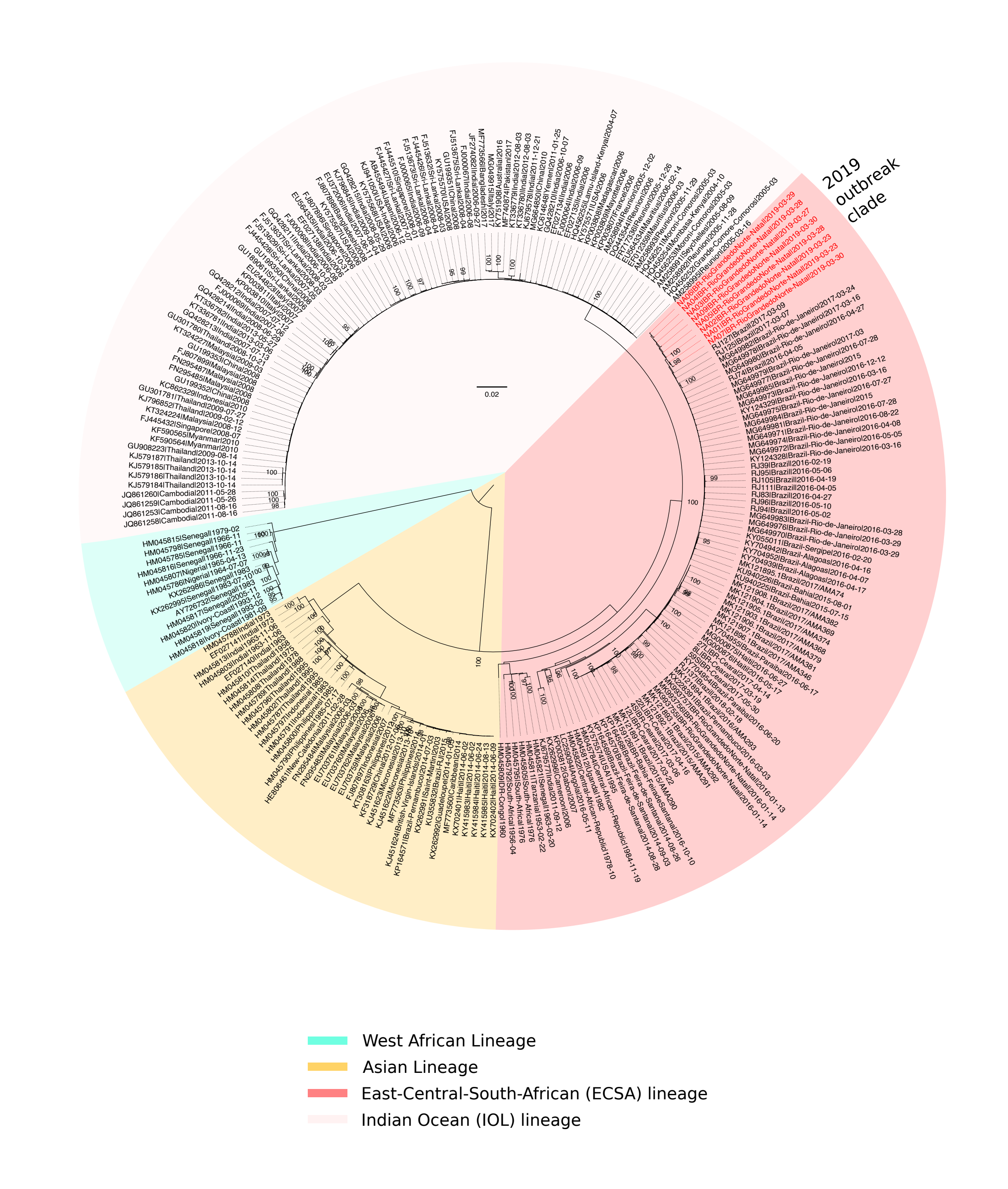
**Figure S1. Global chikungunya phylogeny.** A Maximum Likelihood phylogeny comprising 8 new sequences (coloured in red) generated in this study plus 224 sequences retrieved from NCBI.	Chikungunya virus lineages clades are coloured according to the legend.

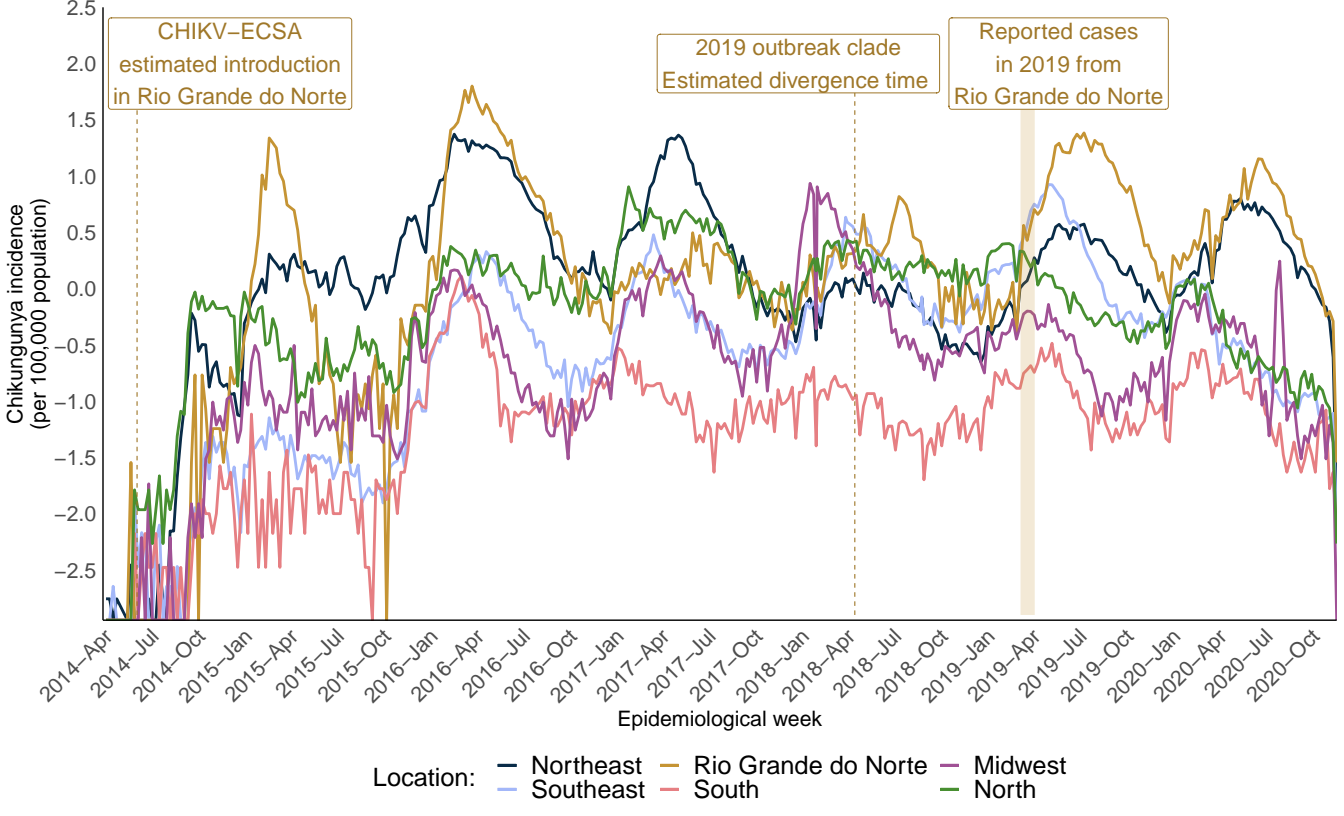
**Figure S2. Time series of chikungunya outbreaks reported in Rio Grande do Norte state and all regions of Brazil.** Epidemic curve of chikungunya virus incidence per 100,000 inhabitants calculated from notified cases from Rio Grande do Norte state and all regions of Brazil. Y-axis values were log-transformed.

**Table S1. Summary statistics of epidemiological and sequencing data.**

| **Sample ID** | **Isolate ID** | **NCBI Accession number** | **Sample type** | **Symptoms onset date** | **Collection date** | **Delay (days) onset and collection date** | **Year of age** | **Sex** | **State** | **Municipality** | **Ct value** | **Reads** | **Coverage (%)** | **Depth of coverage** |
| --- | --- | --- | --- | --- | --- | --- | --- | --- | --- | --- | --- | --- | --- | --- |
| 2746 | NA01 | MW260512 | Serum | 2019-03-21 | 2019-03-23 | 2 | 32 | M | RN | Natal | 21 | 159071 | 93.6 | 9740.8 |
| 2747 | NA02 | MW260513 | Serum | 2019-03-23 | 2019-03-23 | 0 | 19 | M | RN | Natal | 22 | 105093 | 94.5 | 7614.8 |
| 2748 | NA03 | MW260515 | Serum | 2019-03-25 | 2019-03-27 | 2 | 53 | M | RN | Natal | 26 | 280612 | 93.1 | 15545.7 |
| 2749 | NA04 | MW260514 | Serum | 2019-03-26 | 2019-03-28 | 2 | 23 | M | RN | Natal | 34 | 208775 | 92 | 10591.3 |
| 2750 | NA05 | MW260516 | Serum | 2019-03-27 | 2019-03-28 | 1 | 22 | F | RN | Natal | 28 | 147517 | 93.2 | 8447.5 |
| 2751 | NA06 | MW260517 | Serum | 2019-03-24 | 2019-03-29 | 5 | 78 | F | RN | Natal | 29 | 188258 | 92.2 | 10414.6 |
| 2753 | NA07 | MW260518 | Serum | 2019-03-28 | 2019-03-30 | 2 | 78 | F | RN | Natal | 17 | 177192 | 95.2 | 15484.6 |
| 2755 | NA09 | MW260519 | Serum | 2019-03-29 | 2019-03-30 | 1 | 57 | M | RN | Natal | 19 | 223411 | 94 | 11299.2 |
| 2754/2019 | - | - | Serum | 2019-03-27 | 2019-04-01 | 5 | 57 | F | RN | Natal | 34 | - | - | - |
| 2743/2019 | - | - | Serum | 2019-03-20 | 2019-03-23 | 3 | 25 | M | RN | Natal | 37 | - | - | - |
| 2744/2019 | - | - | Serum | 2019-03-19 | 2019-03-23 | 4 | 55 | M | RN | Natal | 36 | - | - | - |
| 2745/2019 | - | - | Serum | 2019-03-21 | 2019-03-23 | 2 | 25 | M | RN | Natal | 33 | - | - | - |
| 2752/2019 | - | - | Serum | 2019-03-28 | 2019-03-30 | 2 | 26 | M | RN | Natal | 30 | - | - | - |

**Table S2. Unique nucleotide substitutions shared by CHIKV isolates from the 2019 outbreak in Natal, Rio Grande do Norte, Brazil.**

| Nucleotide position* | Synonymous substitution | Genes |
| --- | --- | --- |
| 1675 | G→A | nsP1 |
| 2326 | A→ G | nsP2 |
| 3077 | A→ C | nsP2 |
| 3529 | C→ T | nsP2 |
| 5080 | T →C | nsP3 |
| 7869 | C →T | capsid |

*Compared to the NCBI CHIKV reference genome (NC_004162.2) and to isolate KP164568 and MK244641, from Bahia and Rio de Janeiro Brazilian states, respectively.
